# Physical fitness in community dwelling older adults is linked to dietary intake, gut microbiota and metabolomic signatures

**DOI:** 10.1101/793612

**Authors:** Josué L. Castro-Mejía, Bekzod Khakimov, Łukasz Krych, Jacob Bülow, Rasmus L. Bechshøft, Grith Højfeldt, Kenneth H. Mertz, Eva Stahl Garne, Simon R. Schacht, Hajar F. Ahmad, Witold Kot, Lars H. Hansen, Federico J. A. Perez-Cueto, Mads V. Lind, Aske J. Lassen, Inge Tetens, Tenna Jensen, Søren Reitelseder, Astrid P. Jespersen, Lars Holm, Søren B. Engelsen, Dennis S. Nielsen

## Abstract

When humans age, changes in body composition arise along with lifestyle-associated disorders influencing fitness and physical decline. Here we provide a comprehensive view of dietary intake, physical activity, gut microbiota (GM) and host metabolome in relation to physical fitness of 207 community dwelling subjects aged +65 years. Stratification on anthropometric/body-composition/physical-performance measurements (ABPm) variables identified two phenotypes (high/low-fitness) clearly linked to dietary intake, physical activity, GM and host metabolome patterns. Strikingly, despite a higher energy intake high-fitness subjects were characterized by leaner bodies and lower fasting proinsulin-C-peptide/blood glucose levels in a mechanism likely driven by higher dietary-fiber intake, physical activity and increased abundance of Bifidobacteriales and Clostridiales species in GM and associated metabolites (i.e. enterolactone). These factors explained 50.1% of the individual variation in physical fitness. We propose that targeting dietary strategies for modulation of GM and host metabolome interactions may allow establishing therapeutic approaches to delay and possibly revert comorbidities of aging.

## 1. INTRODUCTION

Throughout the course of aging, physical impairment and changes in body composition may arise along with a number of lifestyle-associated disorders influencing physical decline and ultimately frailty (Xue, 2011; Holm et al., 2014). Aging inevitably occurs in all organisms with genetics, epigenetics and environmental exposures (e.g. diet, physical activity) being modulators of the bodily deterioration caused by biological age (Khan et al., 2017). A number of guidelines toward dietary and daily physical activity recommendations are currently available, however, adherence remains a significant challenge (Gopinath et al., 2016). Further, food perception and dietary habits can be strongly altered during the course of life, particularly those traits associated with the loss of appetite (declined senses of smell and taste), occurrence of immune-senescence and deterioration of the gastro-intestinal system (Giezenaar et al., 2016).

During the last decade, the gut microbiota (GM) has been recognized as a signaling hub that integrates dietary habits with genetic and immune signals throughout life (Thaiss et al., 2016; Peters et al., 2019). Many inflammatory and metabolic disorders, such as obesity, diabetes and inflammatory reactions, are linked with GM dysbiosis (Boulangé et al., 2016). Among Irish older subjects frailty has been linked with changing GM signatures (Claesson et al., 2012) and age-related insulin resistance has been found to be regulated by the metabolic activity (e.g. production of short-chain fatty acids – SCFA) of a number of Clostridiales species (e.g. *Clostridium IV*, *Ruminococcus, Saccharofermentans*) and *Akkermansia muciniphila* (Bodogai et al., 2018; Kong et al., 2016; Biagi et al., 2010). Further, low abundance of these bacteria leads to increased leakage of proinflammatory epitopes from the gut to the blood stream (due to leaky gut syndrome) activating monocytes inflammation and subsequently impair insulin signaling in rodents (Bodogai et al., 2018).

It is well-established, that frail older adults are characterized by changed dietary habits and altered GM and metabolic signatures relative to non-frail peers (Claesson et al., 2012; Lustgarten et al., 2014), but whether similar signatures can be identified among non-frail older adults of different physical capacity has, to the best of our knowledge, not been investigated previously. A few studies have focused on frail individuals showing that a reduced consumption of dietary fiber compromises the GM associated production of SCFA required for maintenance of colonic epithelial cells and regulation of immune and inflammatory responses (Kong et al., 2016; Biagi et al., 2010; Claesson et al., 2012). Likewise, GM signatures were found to correspond with frailty-indexes in a large cohort of older adults, whose GM composition were inherently driven by dietary patterns (Claesson et al., 2012). Moreover, metabolites related to GM metabolism (e.g. p-cresol sulfate, indoxyl sulfate), peroxisome proliferator-activated receptors-alpha activation, and insulin resistance likely influence physical function in physically impaired older adults (Lustgarten et al., 2014).

Understanding how dietary intake and physical activity in non-frail older adults alter the GM–metabolome axis, and ultimately the physical fitness and the risk of functional decline, is of great clinical interest for the affected subjects as well as for the society. Furthermore, identifying key components of such multifactorial processes may open opportunities to therapeutically address and possibly treat and prevent the comorbidities of aging (Khan et al., 2017). Based on this framework, we characterized dietary intake, daily physical activity, GM and host metabolome in order to be able to explain physical fitness of non-frail older subjects. To this end, we included 207 individuals (65+ years old, self-supportive and apparently healthy) recruited through the Counteracting Age-related Loss of skeletal Muscle mass (CALM) study (http://calm.ku.dk) (Bechshøft et al., 2016). Our findings demonstrate that physical fitness and function corresponded to signatures of fasting proinsulin and average blood glucose, and characterized by clear differences in energy and dietary fiber intake, daily physical activity as well as differential abundance of GM members and a number of fecal and plasma metabolites.

## 2. RESULTS

### 2.1 Participants inclusion

Two hundred seven individuals were included in this cross-sectional baseline study (Bechshøft et al., 2016). The recruited subjects are population-level representatives of community dwelling, self-supportive and apparently healthy older adults living in the Danish Capital Region with body mass index (BMI) ranging between 18.5 and 37.3 kg/m^2^ (Table 1). Detailed inclusion criteria have been described previously (Bechshøft et al., 2016). From each individual, data were obtained on detailed anthropometric, body-composition and physical-performance measurements (ABPm), average daily physical activity, dietary intake and preferences, GM composition, clinical biomarkers, as well as fecal- and plasma-metabolome adding up to 1,232 analyzed features per subject (Figure S1a).

**Table 1.**
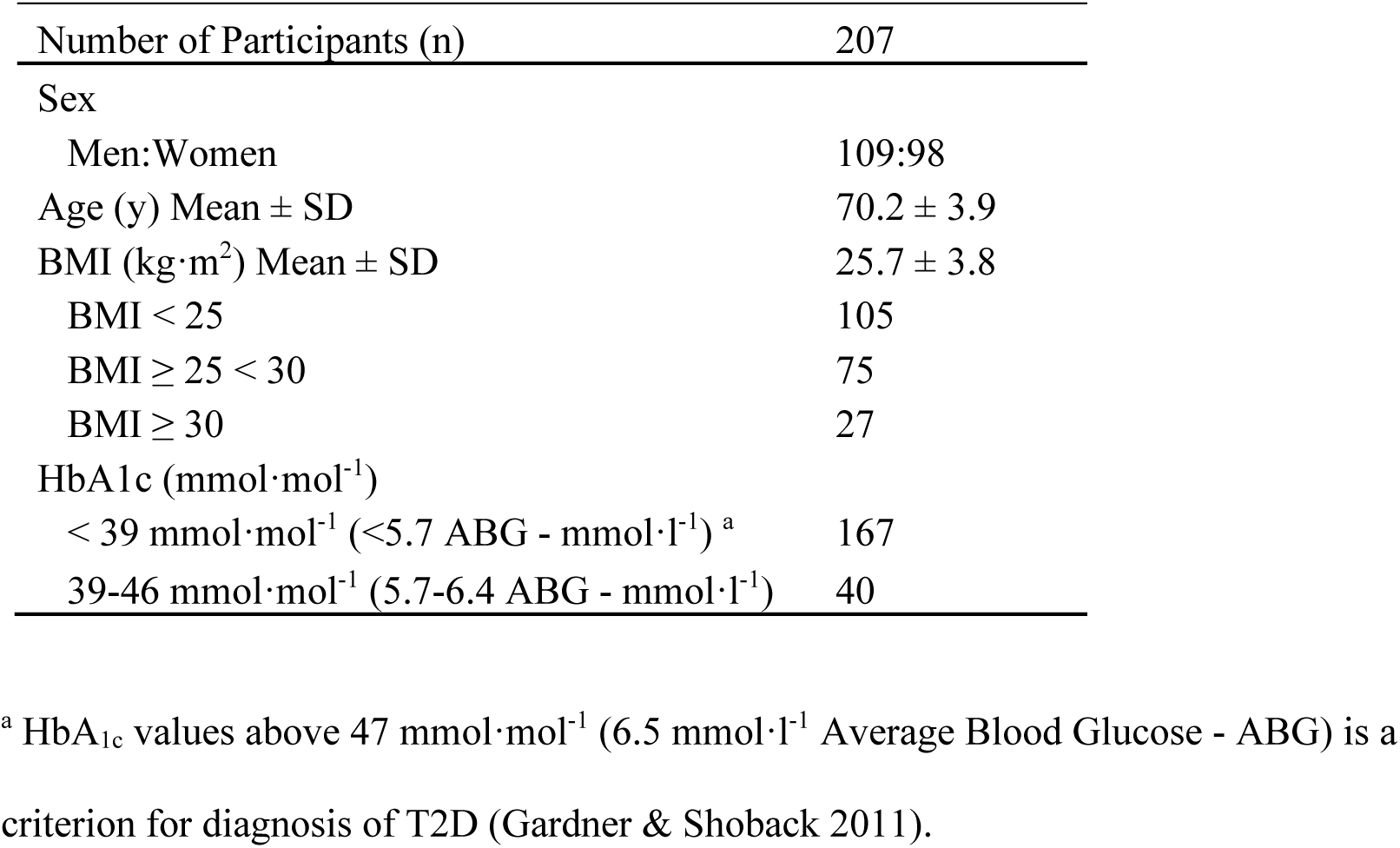
Description of the study participants

### 2.2 Stratification of subjects according to physical fitness and activity monitoring

Study participants were stratified based on non-collinear ABPm variables (Table S1; Variance Inflation Factor, VIF < 2, *r-*coefficient < 0.5) into high- and low-physical fitness phenotypes (level of physical capacity). The selected variables included chair-rise test [30s-test]), BMI, and Dual-energy X-ray Absorptiometry (DXA) scans for body composition (given by legs-soft-tissue fat% (LSF%)), determined as described previously in Bechshøft et al., (2016).

For stratification, hierarchical clustering analysis of principal component analysis (HCP-PCA(Husson et al., 2008)) within sexes was used to determine two fitness phenotypes [high (HF) (n=116) and low (LF) (n=91) (Figure 1a, 1b, Table 2)]. All participants out-performed the suggested ranges for frailty according to the chair-rise test (Jones et al., 1999; Guralnik et al., 1994), while LF phenotypes on average had BMI ranges categorized as overweight (WHO, 2000), as well as a greater deposition of fat mass in their legs (Figure 1b, Table 2). Moreover, 4-day activity monitoring (Dowd et al., 2012) showed significant differences (*p* < 0.001) between the two phenotypes. Longer standing periods (Figure 1c; HF mean: 4.6 ± 1.3, LF mean: 4.2 ± 1.5) and a greater number of steps per day (Figure 1d; HF mean: 11,129 ± 3,861, LF mean: 8,814 ± 3,595) were recorded among HF phenotypes. The habitual daily activity for LF phenotypes was found to be within recommended ranges (taking approximately 7,000-10,000 steps/day (Tudor-Locke et al., 2011)), but the LF subjects were markedly outperformed by the HF subjects (Figure 1d).

**Figure 1.**
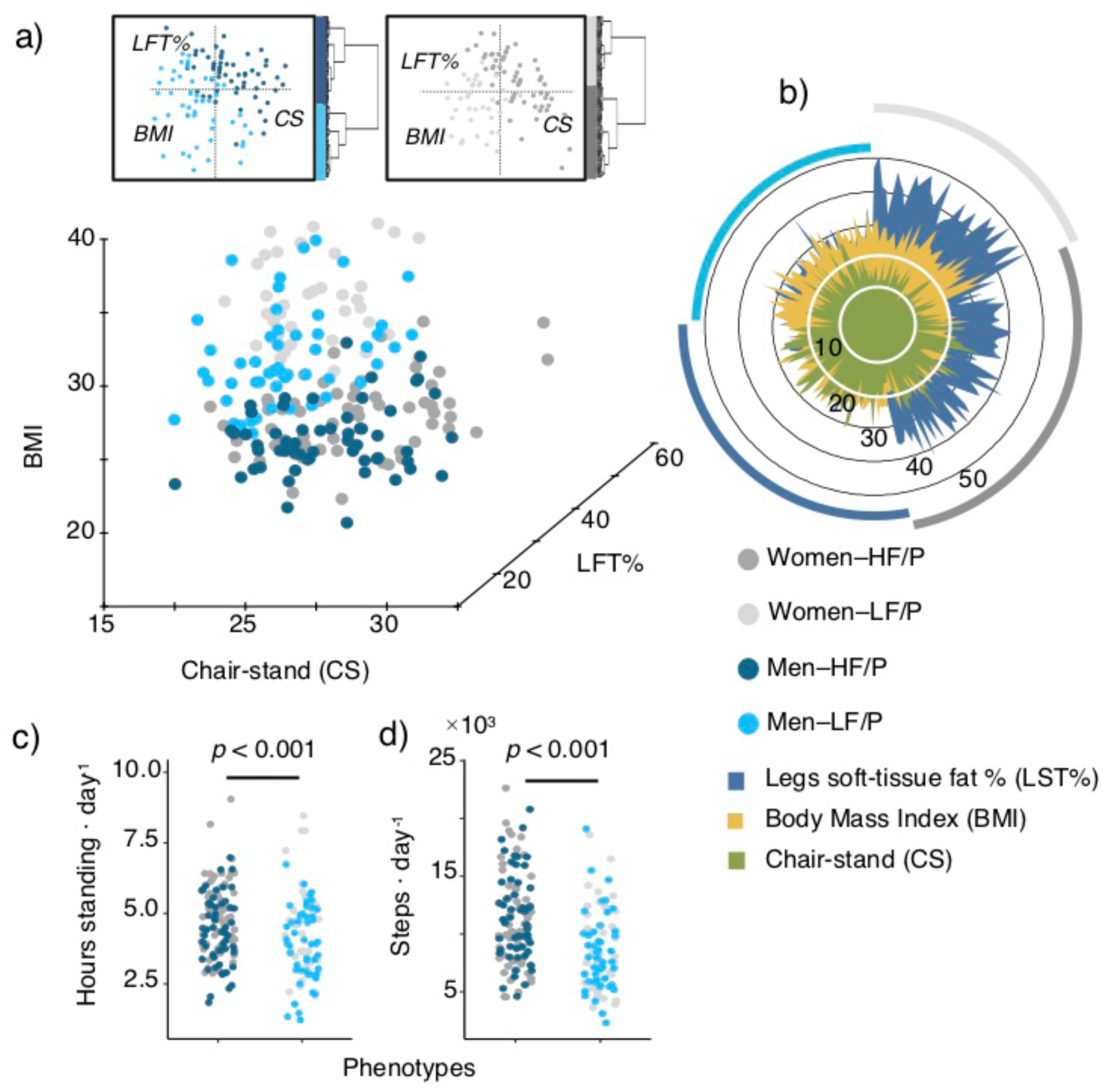
Stratification of fitness phenotypes. (a) Stratification of subjects (n = 207) by hierarchical clustering analysis of principal components analysis (HCA-PCA). Stratification data matrix: [obj x vars] = [207 × 3]. HCA-PCA was performed within sexes and based on ABP measurements. HF/P: high-fitness (n = 116) and LF/P: low-fitness phenotypes (n = 91). (b) ABP measurements distribution among phenotypes and sexes. (c) 4-day activity monitoring displaying hours standing and steps on daily basis for both phenotypes. 4-day activity data matrix: [obj x vars] = [196 × 2]

**Table 2.** Within sex summary of ABP measurements used for stratification of phenotypes (HF/P: high-fitness phenotypes, LF: low-fitness phenotypes).

**Women**
| Functional-Parameter | HF/P | LF/P | *p*-value<sup>a</sup> | Refer. range | Ref. age |
| --- | --- | --- | --- | --- | --- |
| 30s Chair-stand test | 20.6 ± 5.0 | 15.7 ± 3.1 | < 0.001 | 10–16<sup>b</sup> | 65-74y<sup>b</sup> |
| BMI | 22.4 ± 2.1 | 28.9 ± 3.3 | < 0.001 | | |
| LSF% | 35.2 ± 4.0 | 42.7 ± 4.6 | < 0.001 | | |

**Men**
| Functional-Parameter | HF/P | LF/P | *p*-value<sup>a</sup> | Refer. range | Ref. age |
| --- | --- | --- | --- | --- | --- |
| Chair-rise test | 22.9 ± 4.4 | 18.3 ± 3.9 | < 0.001 | 12–18<sup>b</sup> | 65-74y<sup>b</sup> |
| BMI | 24.0 ± 2.2 | 28.3 ± 3.1 | < 0.001 | | |
| LSF% | 20.3 ± 3.4 | 27.0 ± 3.5 | < 0.001 | | |

<sup>a</sup> Comparison between phenotypes was performed by two-tailed Student's *t*-test.664 <sup>b</sup> ref: (Jones et al., 1999; Guralnik et al., 1994)

### 2.3 Dietary food intake in relation to fitness-state

Using 3-day weighted food records (3d-WFR)(Schacht et al., 2019), the daily average energy and macronutrients intake for each person were quantified to obtain an overall view on the dietary intake. On average, the energy intake per person was 24.5 ± 7.4 (range of 11.5 – 55.2) Cal**·**kg body weight^−1^·day^−1^. Protein contributed less of the energy intake (18.9% ± 4.1, range 9-36%) compared to the average energy intake of fat (36.7% ± 7.3, 22-64%) and carbohydrates (44.4% ± 7.7, 17-66%) expressed as percentage of total energy intake.

Total energy consumption per kg body weight (Figure 2a) differed significantly (*p* < 0.001) between phenotypes, with an average daily intake of 29.3 Cal·kg body weight^−1^·day^−1^ in HF phenotypes vs. 23.1 Cal·kg body weight^−1^·day^−1^ in LF phenotypes. The higher energy intake among HF subjects was reflected in a larger fraction of energy (expressed as % energy) from carbohydrates (*p* = 0.01) as compared to that of dietary protein (Figure 2b and Figure S1b). The same pattern was also observed across daily average intake (g·kg body weight^−1^·day^−1^) of dietary fiber (*p* < 0.0001), starch (*p* < 0.0001), simple sugars (*p* = 0.0002) and saturated fatty acids (*p* = 0.0001) (Figure 2c). Moreover, significant (*p* < 0.0001) negative correlations between BMI with dietary fiber consumption (*r* = −0.52) (Figure 2d) energy intake (*r* = −0.52), starch (*r* = −0.35) and simple sugars (*r* = −0.35), as well as positive associations between chair-stand test and energy intake (*r* = 0.25) were found (Figure S1c-f). Questionnaires on food-choices showed that HF subjects to a higher degree than LF subjects consider healthy food as an important element of their daily life (Figure S1g).

**Figure 2.**
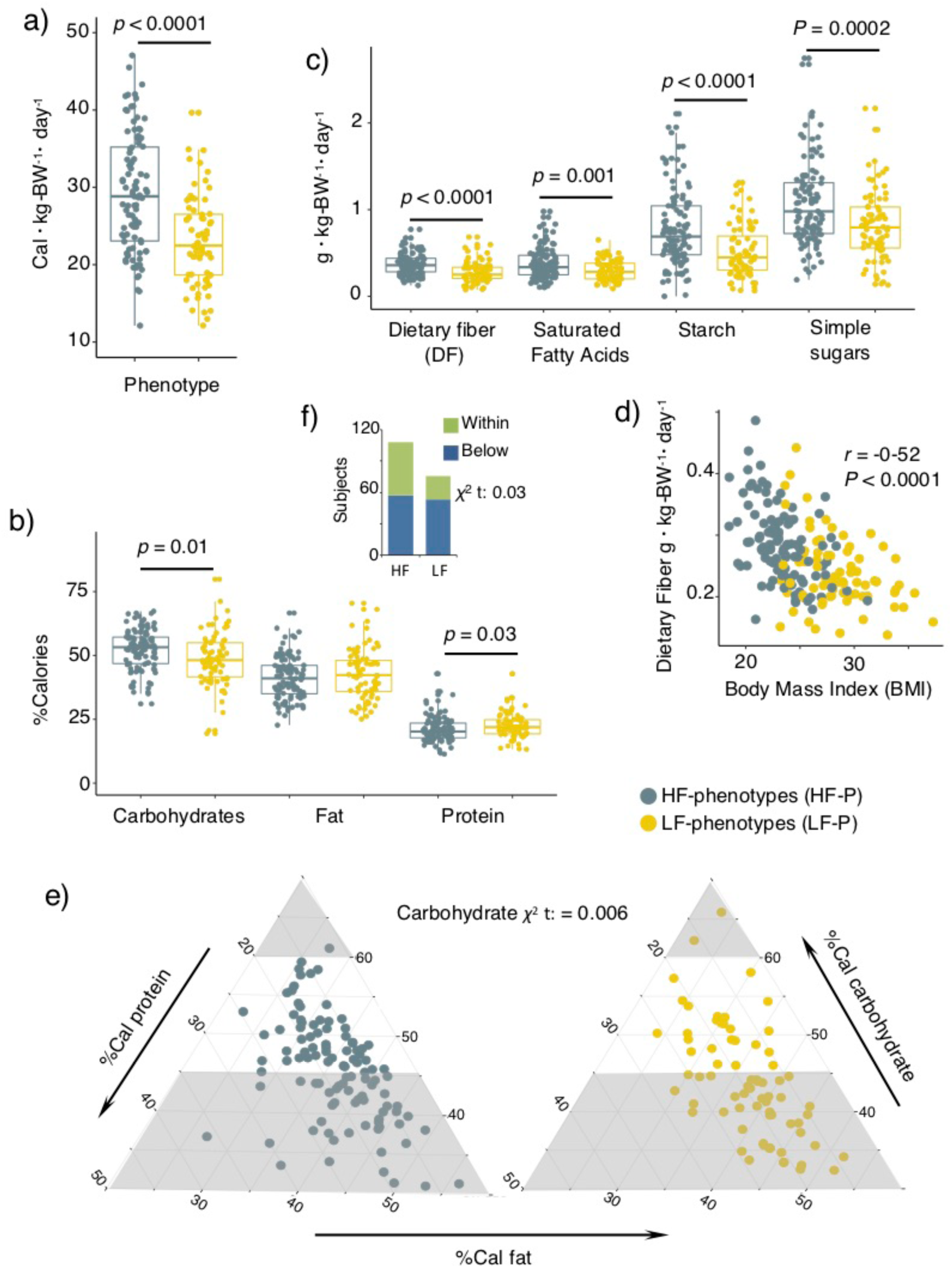
Dietary intake and distribution. (a) Total energy consumption per kg-body-weight per day (Cal·kg-body-weight^−1^·day^−1^) (b) Distribution of Calories proportionally obtained from macronutrients intake in HF and LF phenotypes. (c) Intake of carbohydrates by quality and saturated free fatty acids (g·kg-body-weight^−1^·day^−1^). (d) Pearson correlation between dietary fiber (g·kg-body-weight^−1^·day^−1^) and BMI depicted according to phenotypes category. (e) Proportion of subjects complying with recommended carbohydrates distribution ranges. The gray areas correspond to non-recommended ranges as suggested by the Nordic Nutrition Recommendations. (f) Proportion of subjects complying with recommended distribution ranges of dietary fiber according to the Nordic Nutrition Recommendations. Dietary data matrix: [obj x vars] = [181 × 11]

A considerable proportion of subjects from both phenotypes did not comply with the recommended minimum proportion of energy obtained from carbohydrates (Figure 2e) and dietary fiber intake (Figure 2f) as established by the Nordic Nutrition Recommendations (Nordic Council of Ministers, 2012). Yet, the frequency of compliers-to-non-compliers was significantly higher (carbohydrates: *p* = 0.006, dietary fiber: *p* = 0.03) in HF individuals. Furthermore, using the Goldberg cut-off (Black, 2000), 46 under-reporters (UR) and 2 over-reporters (OR) of energy intake were identified. Nonetheless, if excluded, individuals with higher physical capability (HF phenotype) still had a higher energy (*p* < 0.001) and energy from carbohydrates (*p* < 0.06) intake as compared to LF subjects (Table S2). Since UR and OR subjects did not change the overall findings they were not excluded in downstream analyses.

### 2.4 Characterization of GM and correspondence with fitness and diet

Sequencing of DNA extracted from stool samples yielded 11.3 million reads derived from the 16S rRNA-gene V3-region with an average of 116,476 (48,872 SD) sequences per subject. The analysis of amplicon-sequencing data generated 10,084 zOTUs (sequence variants) summarized over 875 cumulative species (species richness) and 8 core-species (defined as being present in all recruited subjects) among the study subjects (Figure S2a). The relative abundance of core species varied between 18 – 84% (Supplementary Figure 2b). Between sexes no significant differences in beta-diversity (Figure 3a) and alpha-diversity (Figure S2c) were observed. Furthermore, regardless of sex, the study participants were characterized by higher relative abundance of e.g. Lachnospiraceae spp., *Akkermansia* spp., *Blautia* spp., along with reduced proportions of *Bacteroides* spp. (Figure S2d) as compared to the community-dwelling group of older adults recruited for the Irish ELDERMET study (Claesson et al., 2012). This may reflect differences associated with dietary habits, age [mean age: baseline-CALM 70 ± 4y, ELDERMET 78 ± 8y], and geographical location.

**Figure 3.**
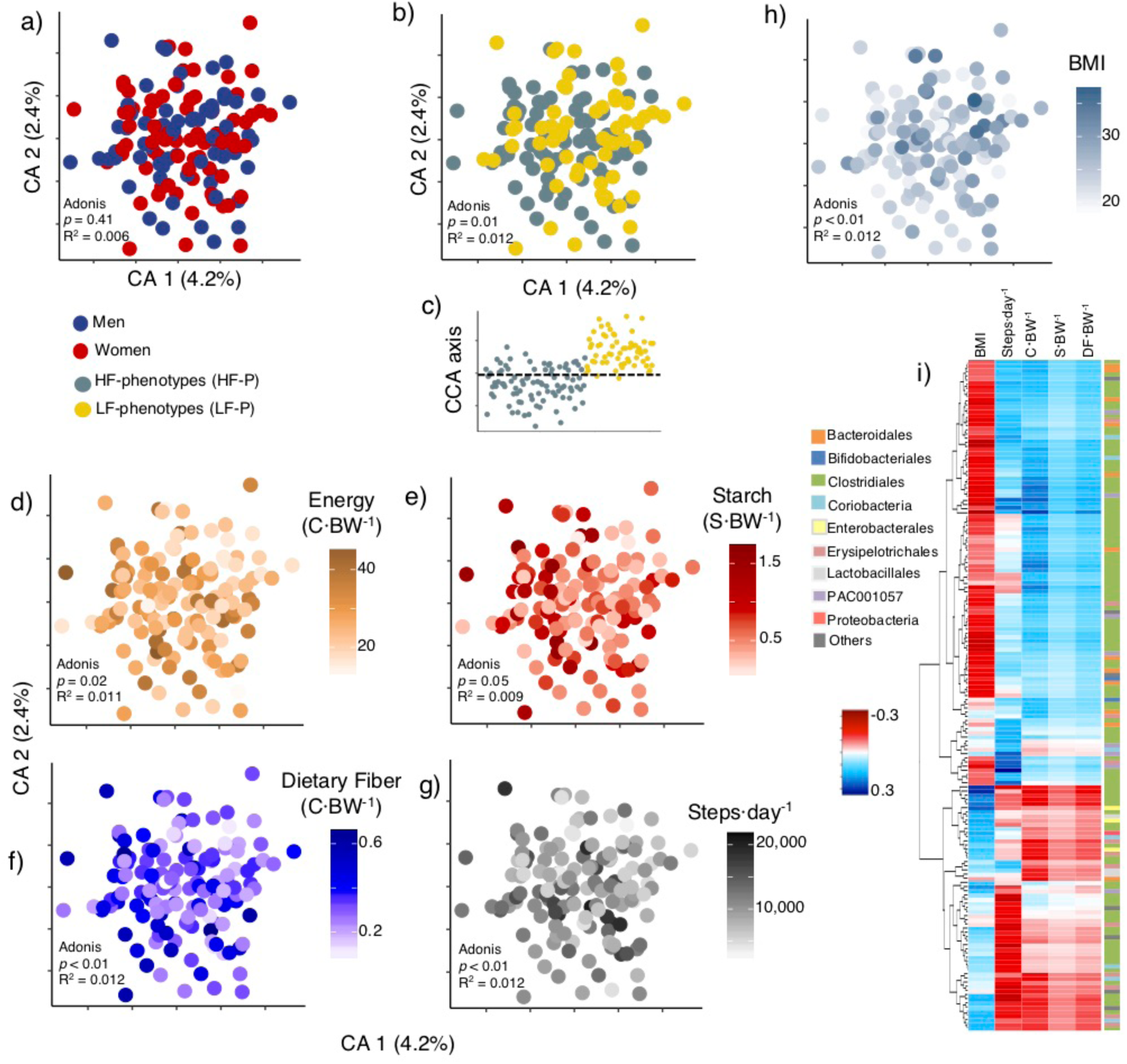
Dietary intake and fitness phenotypes is linked with species-level GM patterns. (a) Gut microbiota (GM) composition determined through Correspondence Analysis of 16S rRNA gene (V3-region) amplicons (summarized zOTUs at species level) determined in the stool samples of the study participants. (b) Correspondence Analysis revealed compositional GM differences between fitness phenotypes. (c) Constrained Correspondence Analysis (CCA) displays discrimination of phenotypes based on permutational test (*p* = 0.03, explained variance = 3.2%). (d) Correspondence Analysis of GM composition depicting gradients of total energy consumption (Cal·kg-body-weight^−1^·day^−1^), intake of (e) starch (g·kg-body-weight^−1^·day^−1^) and (f) dietary fiber (g·kg-body-weight^−1^·day^−1^), (g) steps per day, and (h) BMI. (i) rCC analysis depicting the relationship between gradients of energy consumption, starch and dietary fiber intake, steps per day and BMI, and variations in the abundance of GM members. Heatmap displays the correlation of 161 species with a minimum correlation coefficient of |0.2|*r* from 1^st^ to 3^rd^ components. Species are depicted based on family-level phylogeny. Supplementary Figure 3 displays taxonomy at species level, as well as correlations per canonical axis and explained variance between GM composition and lifestyle covariates derived from rCC analysis. ANOSIM tests were performed on Bray-Curtis distances. GM data matrix: [obj x vars] = [184 × 874]

A substantial higher alpha-diversity (*p* = 0.06, Observed Species) were observed (Figure S2c) among HF phenotypes compared to LF phenotypes, as well as weak but significant (*p* < 0.05) correlations of observed species with BMI, energy and starch intake (Figure S2e-g). Correspondence analysis and analysis of similarities (ANOSIM) on Bray-Curtis (weighted beta-diversity) distance metric calculated from species-level abundance showed significant correspondence (*p* = 0.04) and dissimilarities (*p* = 0.01) in GM composition in connection with the two physical phenotypes (Figure 3b-c).

Also, GM composition was clearly associated with (*p* < 0.05) gradients of energy consumption (Figure 3d), starch (Figure 3e), dietary fiber (Figure 3f) steps per day (Figure 3g) and BMI (Figure 3h) reflecting fitness phenotypes. Using regularized canonical correlation (rCC) analysis associations between those lifestyle covariates (e.g. dietary factors and physical activity) with 161 microbial species were disclosed (Figure 3i, Figure S3) explaining <5% and 13% of the total variance of the microbiota and lifestyle covariates, respectively (Figure S3a). The strongest associations (those > |0.2|*r*, number of species in brackets) were observed for Bacteroidales (12), Bifidobacteriales (2), Clostridiales (106), Coriobacteria (7), Enterobacterales (3), Erysipelotrichales (12), Lactobacillales (3), PAC001057 (Mollicutes members) (8), Proteobacteria (1) and other orders (7) (Figure S3b). Increased intake of energy, starch, dietary fiber, as well as steps per day correlated positively with the relative abundance of up to 103 of those species (e.g. higher Bifidobacteriales abundance) and correlated negatively with BMI (e.g. Proteobacteria being signatures for high BMI) (Figure 3i, Figure S3b).

### 2.5 Host metabolic state in relation to fitness and dietary intake

Untargeted Gas Chromatography-Mass Spectrometry (GC-MS) metabolomics of human fecal extracts and blood plasma as well as targeted SCFA analysis using GC-MS generated a total of 304 analytes (181 analytes in the fecal and 123 analytes in the plasma metabolome). Nearly half of the metabolites variables were identified, either at level 1 or level 2 according to the Metabolomics Standards Initiatives (Sumner et al., 2007). These metabolites were monosaccharides, amino acids, organic acids, sterols and long-, and short chain fatty acids. In addition, 31 biomarkers for immunological function, renal and liver function, as well as glucose and lipid metabolism were acquired through blood clinical profiling.

Correspondence analysis on the combined metabolome blocks showed weak discrimination of sexes (Figure 4a) and pronounced discrimination between fitness phenotype (Figure 4b) based on their metabolic profile. Variations in metabolome composition corresponded clearly (*p* < 0.05) with energy intake and consumption of dietary fiber, starch, simple sugars (Figure 4c-f), as well as steps per day and hours-standing-per-day (Figure 4g-h, including stratifying variables: BMI (Figure 4i), chair-stand and LFT%, Figure S4a-b). Likewise, rCC analysis showed significant associations between lifestyle covariates and 34 clinical/metabolic variables (Figure 4j), explaining 9% and 15% of the total variance of the metabolome and lifestyle covariates, respectively (Figure S4c). The strongest associations (> |0.2|*r*) were observed for 19 clinical biomarkers, 10 gut metabolites and 5 plasma metabolites (Figure 4j). Increased intake of energy, starch, dietary fiber (or dietary covariates), as well as steps per day correlated positively with mono- and di-saccharides and negatively with amino acids (Pro, Ala, Trp), glucose metabolism parameters (proinsulin, glucose HbA1c, HbA1c), lipid metabolism (triglycerides, vLDL) and renal function (creatinine, inversely to estimate glomerular filtration rate (eGFR)) measurements, primary bile acids (lithocholic acid) and N-Nitrosotrimethylurea (Figure 4j). Moreover, a higher proportion of enterolactone in the fecal metabolome of HF subjects were also found (Figure 4k). Remarkably, the concentrations of SCFA as well as other/branched-chain fatty acids (O/B-CFA) in the fecal samples did not differ according to phenotypes (*p* > 0.13) or dietary intake factors (Figure 4l-m).

**Figure 4.**
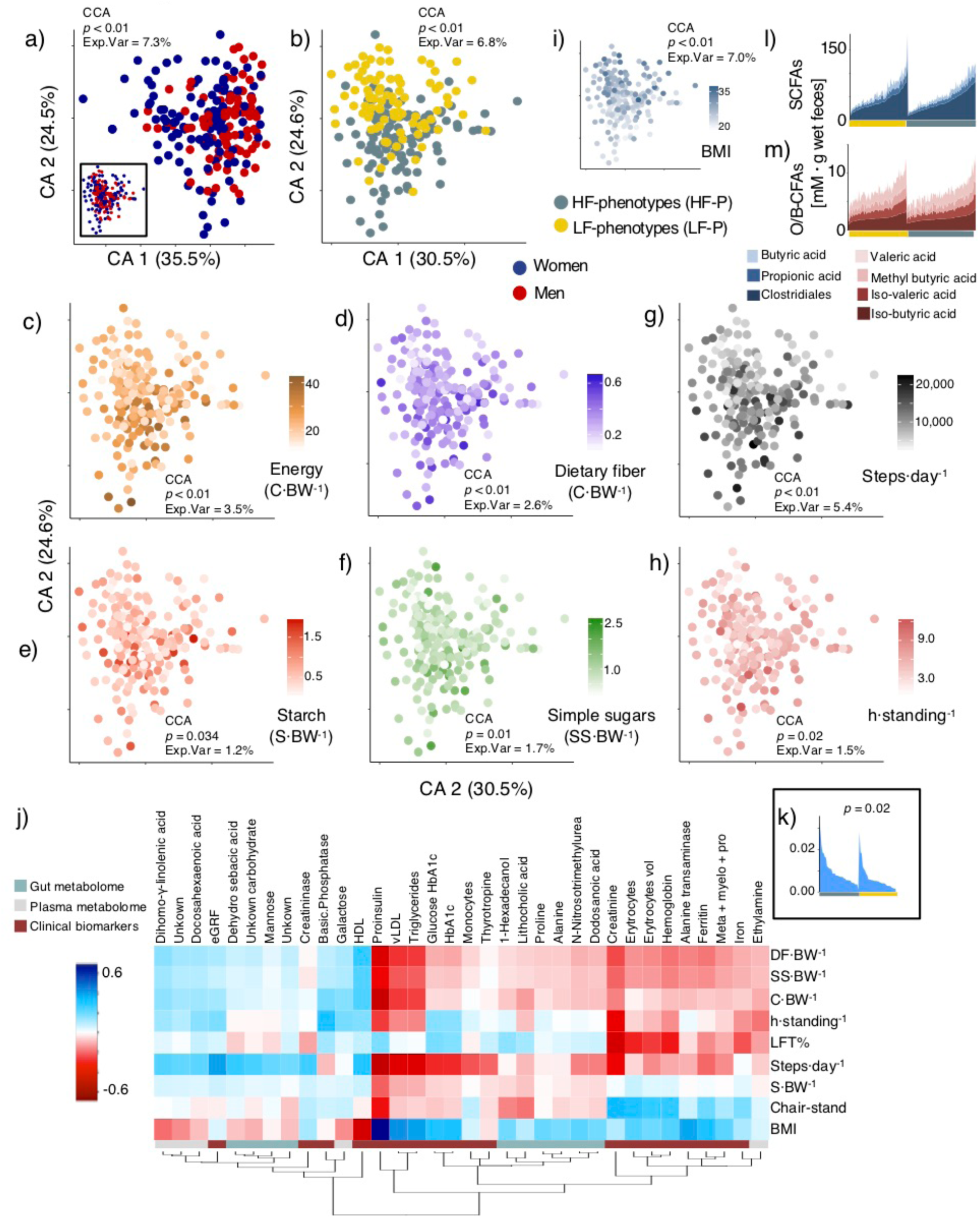
Profiling of host metabolome in relation to dietary intake. (a) Correspondence Analysis on combined fecal-, plasma-metabolomes and clinical biomarkers of the study participants. Significant differences due to sex were determined with constrained correspondence analysis (CCA). Inset shows a partial Correspondence Analysis after conditioning for the cofounding effect of sex. (b) Correspondence Analysis discriminates compositional differences in metabolomic profiles between fitness phenotypes. (c) Correspondence Analysis of metabolites in relation to total energy consumption (Cal·kg-body-weight^−1^·day^−1^), intake of (d) dietary fiber (g·kg-body-weight^−1^·day^−1^), (e) starch (g·kg-body-weight^−1^·day^−1^) and (f) simple sugars (g·kg-body-weight^−1^·day^−1^), (g) steps per day, (h) hours standing, and (i) BMI. (j) rCC analysis showing the relationship between gradients of energy consumption, dietary fiber, starch and simple sugar intake, steps per day, hours standing and BMI, with variations in metabolome composition. Heatmap displays the correlation of 34 clinical/metabolome variables with a minimum correlation coefficient of |0.2|*r* from 1^st^ to 4^th^ components. Supplementary Figure 4 shows correlations per canonical axis as well as explained variance between metabolome composition and lifestyle covariates derived from rCC analysis. (k) Significantly (*t-*test, *p* = 0.02) different relative distributions in enterolactone determined in fecal samples of HF and LF phenotypes (l-m) Range of fecal SCFAs and O/B-CFAs concentrations sorted according to fitness phenotype. Metabolome data matrix: [obj x vars] = [184 × 335]

### 2.6 Dietary intake, gut microbiota and metabolic signatures explain fitness levels independently from physical activity

Characterization of subjects after variable selection based on Random Forest and backward elimination procedure selected 55 variables (Figure 5a) with different levels of importance (Figure 5b) that discriminate the two phenotypes with a high level of accuracy (Figure 5c-d). The features included 25 bacterial species belonging to 7 bacterial orders (Clostridiales, Saccharibacteria, Bacteroidales, PAC001057, Enterobacterales, Erysipelotrichales and Bifidobacteriales), seven dietary components (energy, saturated fatty acids, simple sugars, starch and dietary fiber intake, and energy derived from proteins and carbohydrates); five clinical biomarkers (alanine transaminase, triglycerides, vLDL, fasting proinsulin, average blood glucose/HbA1c). In addition, seven plasma metabolites (amino acids and organic acids), ten fecal metabolites (sugar alcohols, amino acids, primary bile acids and urea) and physical activity (steps per day) were also tabbed (Figure 5a).

**Figure 5.**
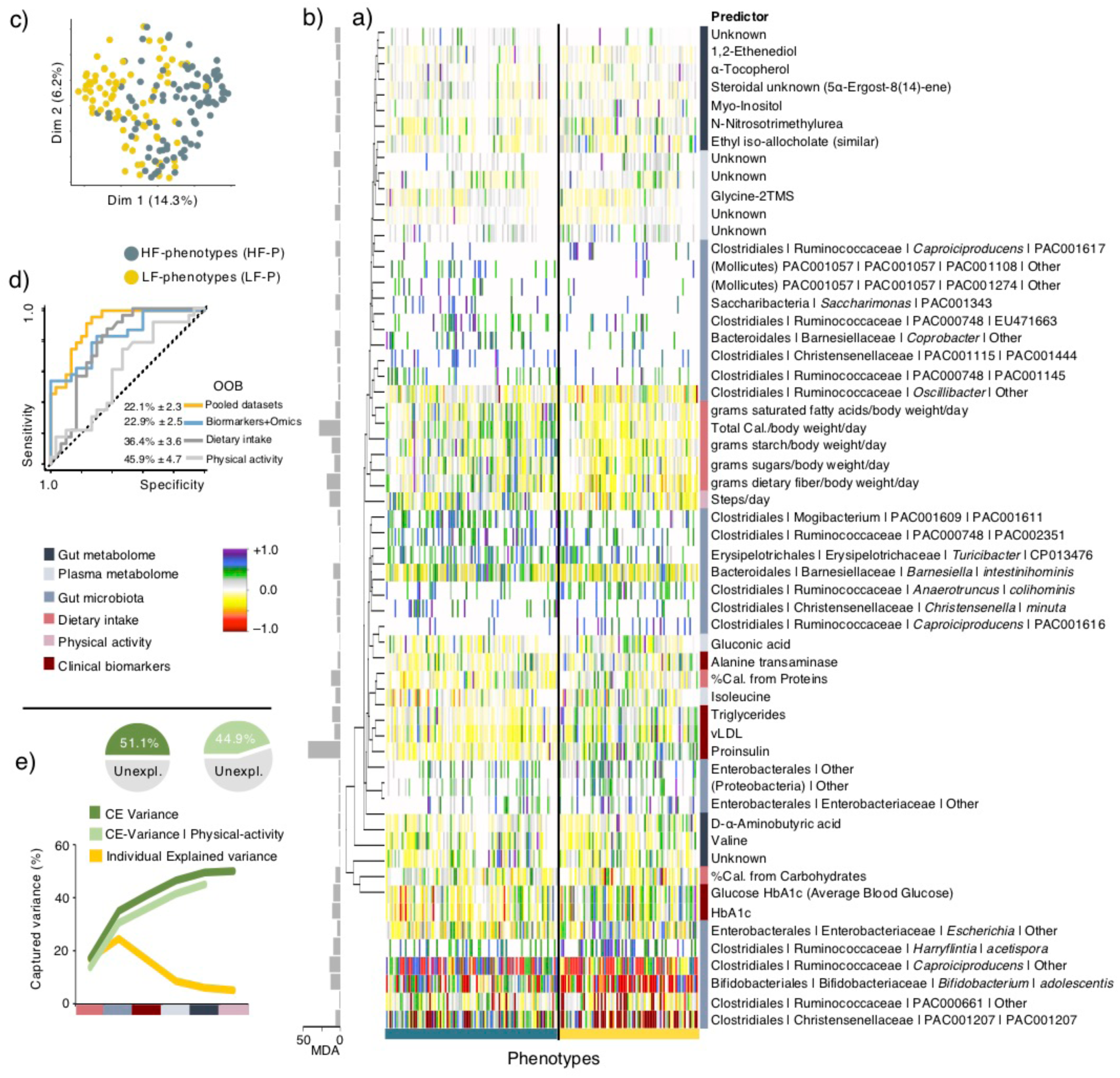
Signatures discriminating physical phenotypes. (a) Heatmap displaying mean centered normalized abundance of 56 features selected using Random Forest towards discrimination of phenotypes and (b) their importance as determined on the basis of Mean Decrease in Accuracy. (c) Multidimensional scaling plot discriminates subjects’ phenotype based on the selected features. (d) ROC curves and out-of-bag error rate (OOB) for Random Forest classifier based on the selected variables, for combined datasets (all selected features), GM and metabolome, dietary intake and physical activity (e) Captured variance for fitness variables (BMI, chair-stand and LSF%) as a function of selected features through redundancy analysis (RDA). Individual Explained Variance displays the size effect of a given dataset, CE-Variance represents the cumulative explained variance and CE-variance | physical-activity shows the accumulative explained variance conditioned by physical activity. Pie charts summarize the total proportion of explained variance before and after conditioning for physical activity. Data matrix: [obj x vars] = [181 × 56]

Discrimination of the two phenotypes based on all the selected features (combined datasets) had the highest level of accuracy (22% out-of-bag error rate, OOB), followed GM and clinical/metabolome features (23% OOB), dietary intake (36% OOB) and physical activity parameters (46% OOB) (Figure 5d). Through redundancy analysis (RDA) the effect of the selected variables (within blocks) on the stratifying variables it was found that GM had the largest explanatory power (24.7%), followed by dietary intake (17.3%), clinical biomarkers (16.8%), gut metabolome (8.8%), plasma metabolome (6.2%) and physical activity (5.2%) (Figure 5e). Notably, the cumulative explained variance conferred by the pool of selected features reached 50.1%, and even after conditioning the effect of physical activity over the stratifying variables, the cumulative explained variance reached up to 44.9% (Figure 5f).

## 3. DISCUSSION

The number of older-adults over the age of 65 will increase by more than 50% worldwide over the next three decades (NIH 2011), potentially with huge implications for the health and economy of the implicated individuals and society as a whole. With this, understanding the physical mechanisms and lifestyle conditions linked to fitness and independence in older adults becomes a relevant field of research.

Despite the homogeneity of the recruited subjects (all non-frail and without serious disease) noticeable differences in fitness level was observed and based on non-collinear ABPm variables (chair-rise test, BMI and DXA-scan based body composition), two fitness phenotypes (LF and HF) were identified. Neither of the fitness types were frail (Guralnik et al., 1994), nevertheless dietary, GM and host metabolome factors were found to clearly discriminate between the two fitness types. HF subjects were characterized by a higher consumption of foods of plant origin as also reflected by their higher levels of total carbohydrates (i.e. starch, simple sugars) and dietary fiber, accompanied by a higher adherence to the recommended intake of carbohydrates and dietary fiber intake given by the Nordic Nutrition Recommendations (Nordic Council of Ministers 2012). These differences were observed in spite of the methodological limitations of 3d-WFR to capturing long term variability (Yang et al., 2010). Furthermore, whether awareness of dietary guidelines influenced the selection of dietary choices in the study participants remains to be investigated, but it is worth mentioning that HF subjects consider healthy food as an important component in their life as also described by Schacht et al., (Schacht et al., 2019).

The GM community and host metabolome clearly discriminated between the HF and LF phenotypes and was largely associated with the consumption of total energy, and plant derived nutrients (such as starch and dietary fibers as well as enterolactone, all being higher in HF subjects). A number of features (Figure 5a) selected from GM, host metabolome, dietary intake and daily physical activity were able to strongly discriminate and explain variation between phenotypes, thereby indicating their strong association with physical function. Daily physical activity showed the lowest power towards phenotypic differentiation (in spite of the high validity of the method for activity monitoring (Dowd et al., 2012)) and explaining only 5% of the phenotypic variance. Albeit conditioning for physical activity, the remaining set of selected features explained up to 45% of the total variance of the stratifying variables. In particular dietary intake (17% of explained variance), GM composition (24%) and host metabolome (25%) signatures are important drivers of phenotypic differentiation (Figure 5), and also described in animal models (Fujisaka et al., 2018). Accordingly, HF subjects showed a higher proportion of GM members commonly known for their protective roles, such as *Bifidobacterium adolescentis* and *Christensenella* species (Goodrich et al., 2014), and whose abundance corresponded negatively with glucose and lipid metabolism biomarkers (proinsulin, HbA1c, vLDL, triglycerides). Contrarily, LF phenotypes had increased levels of the same biomarkers and a higher relative abundance of pro-inflammatory microbial members in the gut, as for example Enterobacterales (Fei & Zhao 2013; Hoarau et al., 2016; Khan et al., 2014).

SCFAs derived from GM activity have been identified as signaling molecules responsible for maintenance of the integrity of colonic epithelium, glucose homeostasis, lipid metabolism and appetite regulation (Morrison et al., 2016). Claesson et al., (Claesson et al., 2012) reported higher SCFA concentrations (acetate, butyrate and propionate) in the fecal metabolome of older adults living as community-dwellers compared to frail individuals living in residential care. Moreover, decreasing concentrations of these SCFAs were associated with advanced levels of frailty given by diet and specific transitions in GM composition (Claesson et al., 2012). However, in the present study no correlations between fecal SCFA and O/B-CFA concentrations with neither macronutrient distribution or fitness phenotype were found. This suggest that levels of physical function amidst healthy older adults may not be primarily dependent upon changes in the production of these compounds. Instead, this could be due to signals of glucose metabolism deterioration as reflected by significantly (*p* < 0.001) higher proinsulin levels and higher average blood glucose (determined by HbA1c-levels) in the LF phenotypes (1/116 HF and 20/91 LF subjects had higher than normal ranges of proinsulin (Chi-Squared *p* < 0.001), 10/116 HF and 30/91 LF had higher ranges than those recommended for HbA1c (Gardner & Shoback 2011) (Chi-Squared *p* < 0.001), see Table S3). High concentrations of proinsulin indicates high insulin secretion and hence diminished peripheral insulin sensitivity resulting in a number of metabolic conditions, compromising muscle strength and physical performance (Segerström et al., 2011). Proinsulin was the most important feature of phenotype discrimination and corresponded inversely with the abundance of *Bifidobacterium adolescentis* and several species of *Christensenella*, and Ruminococcaceae (Figure 5a), strongly indicating that GM-proinsulin interactions could be mediators of fitness phenotype. *Bifidobacterium* species (including *B. adolescentis*) have previously been described as promoters of adiponectin and decreasing expression of interleukin-6, both playing prominent roles in metabolic derangements associated with glucose regulation and fatty acid oxidation (Su et al., 2015; Straub & Scherer 2019; Aoki et al., 2017). *Christensenella minuta* (another Clostridiales member) is enriched in individuals with low BMI and has been demonstrated to reduce weight gain and adiposity in mice (Goodrich et al., 2014). Furthermore, while playing a protective role against inflammation, some Clostridiales members act as promoters of regulatory T-cells by interacting with toll-like receptors 2 (TLR2) on intestinal epithelial cells (Kashiwagi et al., 2015). Contrarily, species of Enterobacterales have been consistently linked with insulin resistance and inflammatory responses (Fei & Zhao 2013; Hoarau et al., 2016; Khan et al., 2014), and by means of cell epitopes (i.e. LPS) they interact with TLRs triggering pathogen recognition, low-grade inflammation (Franceschi & Campisi, 2014) and fat accumulation in adipose tissue that ultimately influence muscle strength (Boulangé et al., 2016).

In summary, our findings suggest that dietary patterns underlie mechanisms of physical phenotype differentiation among well-functioning community dwelling older adults, particularly as a driver of GM and glucose metabolism interactions. Despite the limitations of this study related to its inherent cross-sectional nature, the results provide strong evidence emphasizing the central role of diet towards the onset of physical deterioration and its implications prior to clinical manifestations of frailty, e.g. muscle composition and diminished strength (Xue, 2011). Many of the dietary, GM and metabolomic signatures seen in frail older adults (Claesson et al., 2012; Bodogai et al., 2018; Kong et al., 2016; Lustgarten et al., 2014) are already evident in the non-frail, community-dwelling older-adults of low-fitness of this study, pointing at the importance of early intervention strategies, also in this age group. Thus, in view of these findings, developing strategies to improve awareness and adherence to dietary recommendations (complying with dietary reference intakes or even with personalized nutrition (Zeevi et al., 2015)), targeting the regulation of GM and host metabolome interactions, can open opportunities to delay the comorbidities of aging.

## 4. EXPERIMENTAL PROCEDURES

### 4.1 Study Participants

Procedures of the CALM project (Clinical Trials NCT02115698) were approved by the Danish Regional Committees of the Capital Region (H-4-2013-070), performed according to the Declaration of Helsinki II and the experimental designed followed as previously described (Bechshøft et al., 2016). For the current study, two hundred and seven subjects (65+ years of age) were selected at baseline of the CALM intervention project following the criteria described in Bechshøft et al.,(Bechshøft et al., 2016). Participants were not allowed to take part in any organized sports or resistance training more than once a week, did not suffer from defined metabolic-, tissue-, or gastro-intestinal disorders, nor were prescribed antibiotics 3 months prior sample collection and enrollment.

### 4.2 Samples and metadata collection

At baseline, participants completed a 3-day weighted food record where food and beverage intake were registered for 3-consecutive days (Wednesday to Friday). Dietary information was typed into the electronic dietary assessment tool, VITAKOST™ (MADLOG APS, Kolding, Denmark), which uses the Danish Food Composition Databank (version 7.01; Søborg; Denmark) to estimate individual energy and macronutrient intake.

Fecal and blood plasma samples were collected and handled according to the following procedures: (i) fecal samples were kept at 4°C for maximum 48 h after voidance, and stored at −60°C until further use; (ii) overnight-fasted-state (OFS) plasma-samples were collected and deposited in heparin, centrifuged at 3,000×g for 10 min at 4°C, and then stored at −60°C.

For screening of blood-biomarkers, the following tests were performed: complete blood count (CBC), proinsulin-C-peptide (P-CP), glycosylated hemoglobin (HbA1c), coagulation factor, estimate glomerular filtration rate (eGFR), thyroid-stimulating hormone (TSH), and iron-ferritin test determined as described in Bechshøft et al., (Bechshøft et al., 2016) For anthropometric and functional capacities, height (cm) and body-weight (kg) in OFS were measured. Average fast-pace gait speed was measured on an indoor 400 m horizontal track. Number of chair-stands in 30s from a standard table chair was recorded. Relative legs-soft-tissue fat% (LSF%) was determined as an estimate of legs-soft-tissue fat-free and fat-mass based on a dual energy x-ray absorptiometry (DXA) scan (Lunar iDXA Forma with enCORE Software Platform version 15, GE Medical Systems Ultrasound & Primary Care Diagnostics, Madison, WI, USA) performed on participants in overnight fasted state.

### 4.3 Quantitative questionnaires on food habits

Quantitative questionnaires contained information on food habits, perceptions and preferences, as well as information about life style changes and dietary habits over the life course (Bechshøft et al., 2016).

### 4.4 GM and metabolomics

Procedures for profiling and process GM and metabolomics data are described in Supplementary Methods.

### 4.5 Statistical Analyses

Stratification of individuals was based on ABP measurements using the variables described in Table S1. Collinear variables were initially removed, leaving chair-stand [30s-test]), DXA scans (legs-soft-tissue fat% determined in both legs) and BMI as features with a variance inflation factor (VIF) < 2 and *r-*coefficient < 0.5. Subjects were divided according to sex, and a hierarchical clustering analysis of principal component analysis (Husson et al., 2008) was performed on the selected variables (100 iterations).

For univariate data analyses, pairwise comparisons were carried out with unpaired two-tailed Student’s *t*-test, Pearson’s coefficient was used for determining correlations and Chi-Square test for evaluating groups distributions. For multivariate data analyses, the influence of covariates (e.g. dietary components, BMI, etc.) on data blocks (GM and metabolome) were assessed with (Constrained-) Correspondence Analysis with permutation tests (1,000 permutations), as well as analysis of similarities (ANOSIM test, 999 permutations) on Bray-Curtis distances (implemented in the *Vegan* R-package (Oksanen et al., 2015)).

Correlation of covariates with the same datasets were determined with regularized canonical correlation (rCC) analysis using the *mixOmics* R-package (González et al., 2012). rCC was crossed-validated (leave-one-out approach) with grids (lambda 1 & 2) of 0.05 to 1.0 and a length of 20.

Feature selection for combined datasets was performed with Random Forest. Dataset was randomly divided 200x (200 subsets) into training (70%) and test sets (30%), keeping this proportion over the number of subjects within each fitness group for every split. For a given training set, the *party* R-package (Hothorn et al., 2016) was run for feature selection using unbiased-trees (cforest_unbiased with 6,000 trees) and AUC-based variable (varimpAUC with 100 permutations), and subsequently the selected variables were used to predict (6,000 trees with 1,000 permutations) their corresponding test set using *randomForest* R-package (Liaw & Wiener 2014). The features derived from the subset with a prediction rate within 1 SD above the mean-prediction (based on the 200 subsets) were selected and subsequently, subjected to sequential rounds of feature selection (following the same tuning of unbiased-trees and AUC-based variable) until prediction could no longer improved. Variation partitioning of stratifying variables (BMI, CS and LSF%) based on selected features derived from the different datasets (i.e. GM, diet, host-metabolome, physical activity) was performed using redundancy analysis (RDA) (Oksanen et al., 2015).

## ACKNOWLEDGEMENTS

This project was supported by the University of Copenhagen-funded project “Counteracting Age-related Loss of Skeletal Muscle (CALM)”, the Innovation Foundation Denmark-funded project COUNTERSTRIKE (4105-00015B), stipends from the University of Copenhagen, as well as a PhD-stipend from Universiti Malaysia Pahang, Malaysia and Ministry of Education, Malaysia.

## AUTHOR CONTRIBUTIONS

Conceptualization: D.S.N., J.L.C., S.B.E., L.H., A.P.J.; Methodology: D.S.N., J.L.C., S.B.E., L.H., A.P.J., A.J.L., T.J., S.R., R.L.B.; Formal Analysis: J.L.C., B.K., Ł.K., D.S.N., S.B.E., L.H; Writing – Original Draft: J.L.C., D.S.N.; Investigation, review & editing: all authors; Visualization: J.L.C., D.S.N.; Supervision: D.S.N., S.B.E., L.H.; Funding Acquisition: D.S.N., S.B.E., L.H., A.P.J.

## Data Availability

Sequence data is available at the European Nucleotide Archive, accession number ENA: PRJEB33008 ([dataset] Castro-Mejía et al., 2019). The remaining data that support the findings of this study are available on request from the corresponding authors. The data are not publicly available due to privacy or ethical restrictions.

## SUPPORTING INFORMATION

Additional supporting information can be found in supplementary files.

- Figure S1. Data overview and dietary intake
- Figure S2. GM overview, cumulative- and core-species
- Figure S3. rCC analysis between GM and lifestyle components
- Figure S4. Metabolome correspondence and correlation
- Table S1. Subjects Stratification
- Table S2. Dietary evaluation
- Table S3. Proinsulin and HbA1c levels
- Supplementary Methods

---

**Supplementary Fig. 1.**
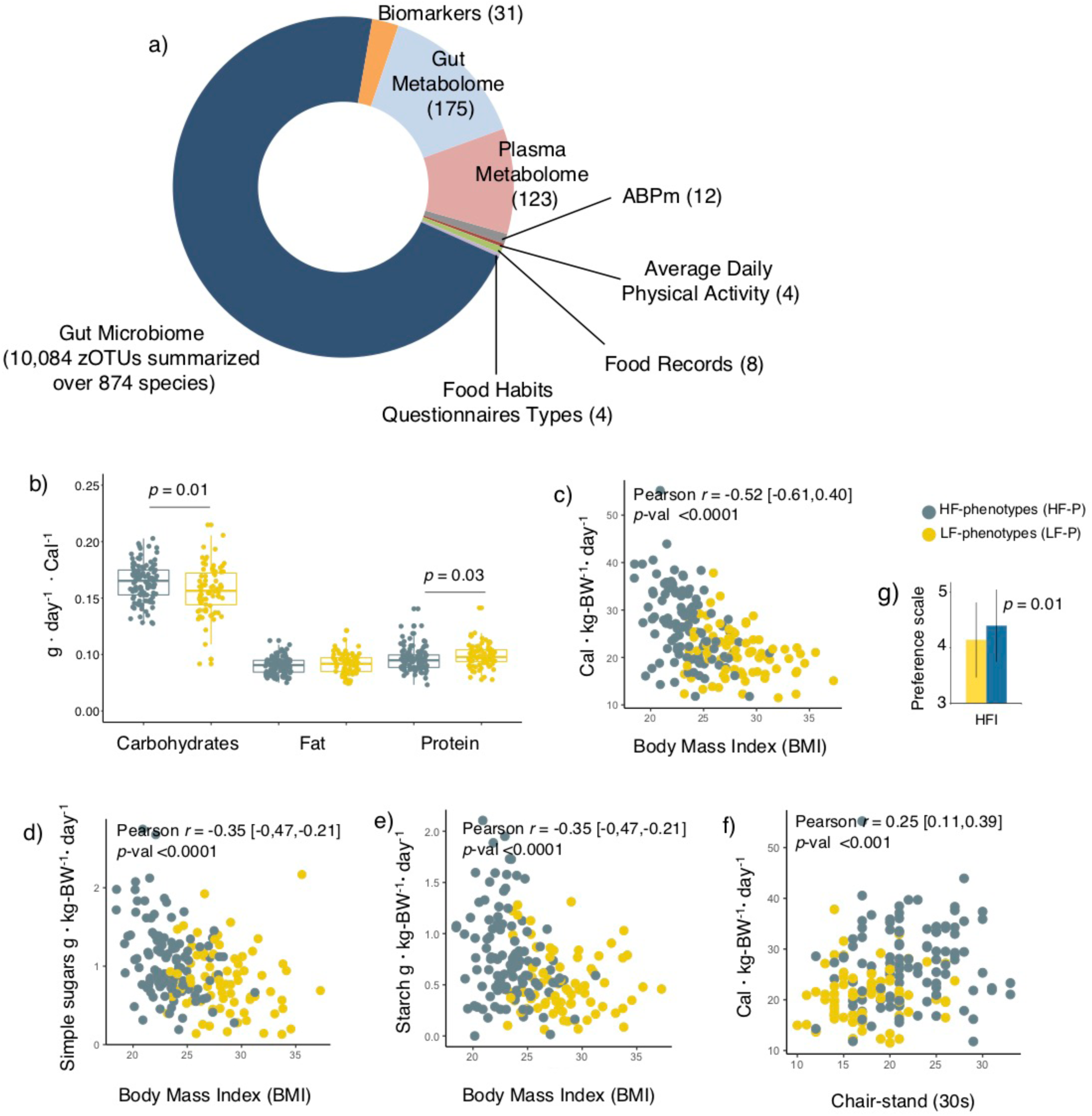
Subjects characterization and dietary intake. (a) Ring chart displays the proportion of variables used for every category at which individuals were characterized (b) Distribution of daily macronutrient intake (g day^−1^) normalized by the total energy intake (Cal) (c) Correlation between energy intake (Cal · kg-Bw^−1^· day^−1^) *vs* BMI (d) Correlation between starch intake (g · kg-BW^−1^· day^−1^) *vs* BMI (e) Correlation between simple sugars intake (g · kg-Bw^−1^· day^−1^) *vs* BMI (f) Correlation between energy intake (Cal · kg-BW^−1^· day^−1^) *vs* Chair-stand test (g) Degree of agreement for food-choices questionnaires: healthy food in an important element of everyday life (HFI). This was evaluated on a scale of 1 – 5. I: is not important 5: is very important,

**Supplementary Fig. 2.**
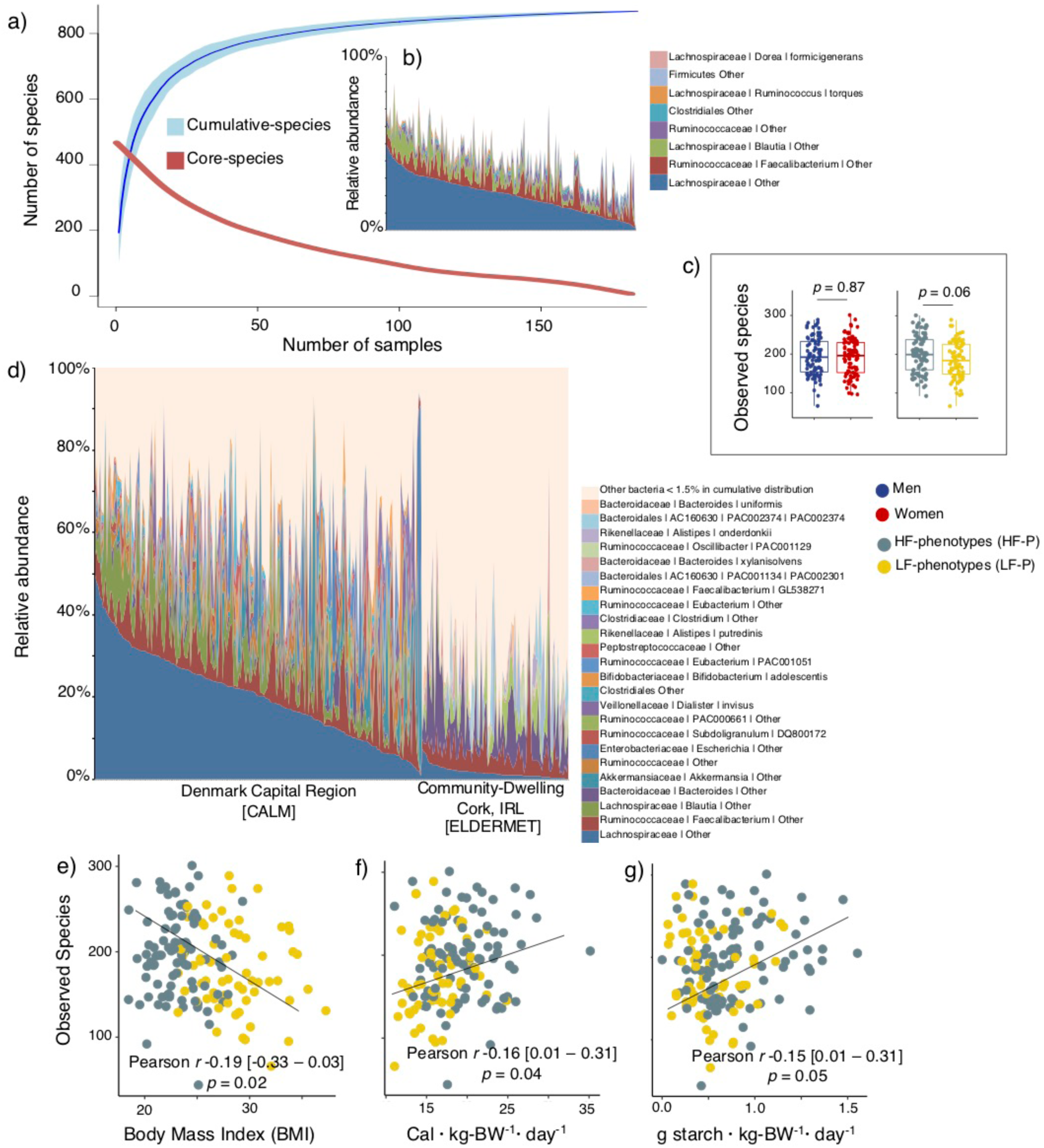
GM overview, cumulative- and core-species. (a) Cumulative- and core species comprised in the CALM study with increasing number of subjects. (b) Relative abundance of core-species (phylotypes summarized to species level). (c) Alpha diversity (of summarized zOTUs at species level) between sexes (*p* = 0.42) and phenotypes (*p* = 0.04) determined by Monte Carlo permutation (100) test. (d) Distribution of species across the subjects of the CALM intervention (16S rRNA v3 region) and the community-dwellers of the ELDERLMET study (16S rRNA v4 region, 454 pyro-sequencing). The publically available sequencing data from the ELDERMENT study were retrieved and analyzed using the parameters described in methods. (e) Correlation between observed species *vs* BMI (f) Correlation between observed species *vs* Energy intake (Cal · kg-BW^−1^· day^−1^) (g) Correlation between observed species *vs* Simple sugars intake (g· kg-BW^−1^· day^−1^)

**Supplementary Fig. 3.**
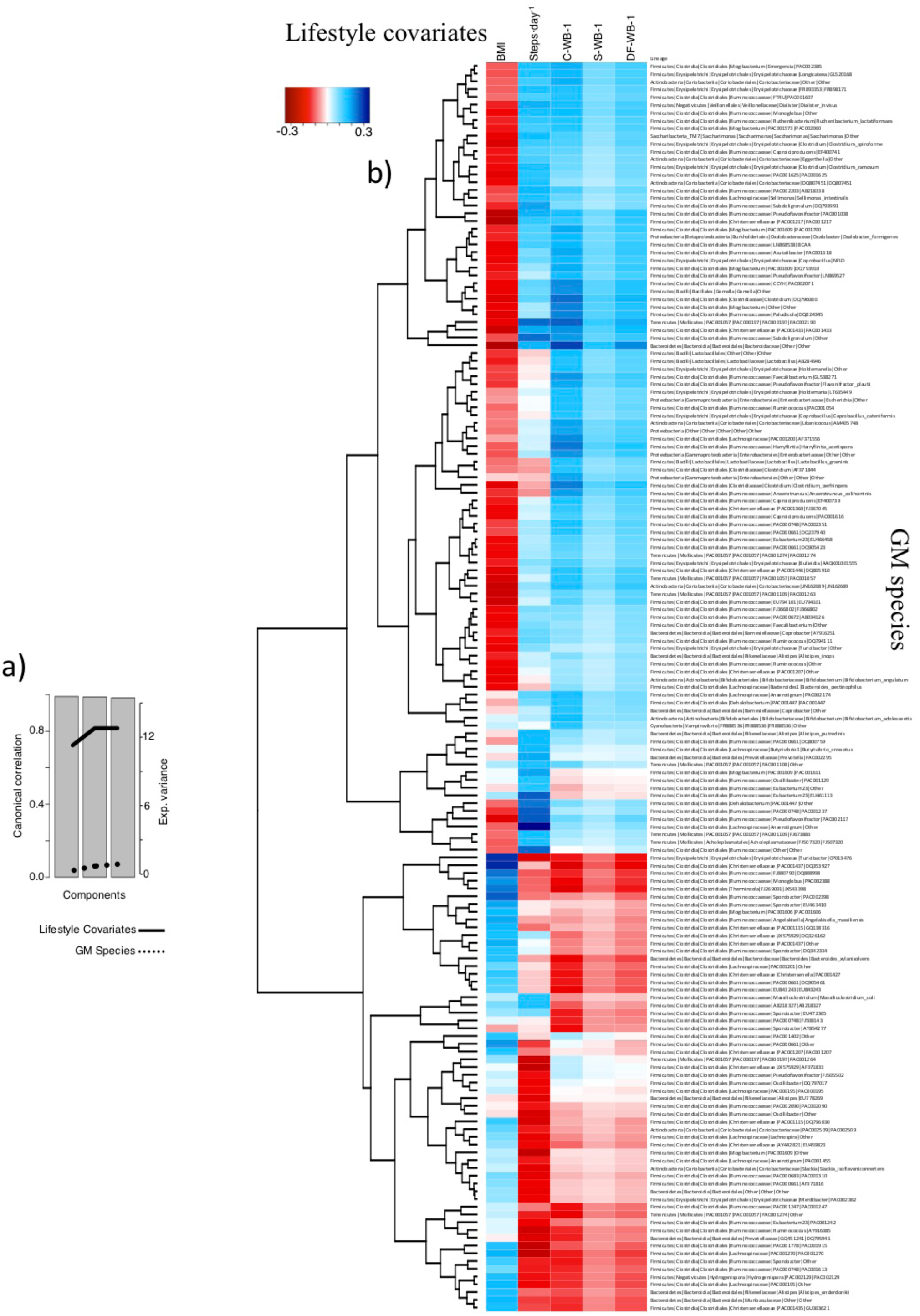
rCC analysis between GM and lifestyle components. (a) Canonical correlation within 1^st^ to 3^rd^ components as well as explained variance between GM composition and lifestyle covariates. (b) Associations between GM species and lifestyle covariates with a minimum correlation coefficient of |0.2|

**Supplementary Fig. 4.**
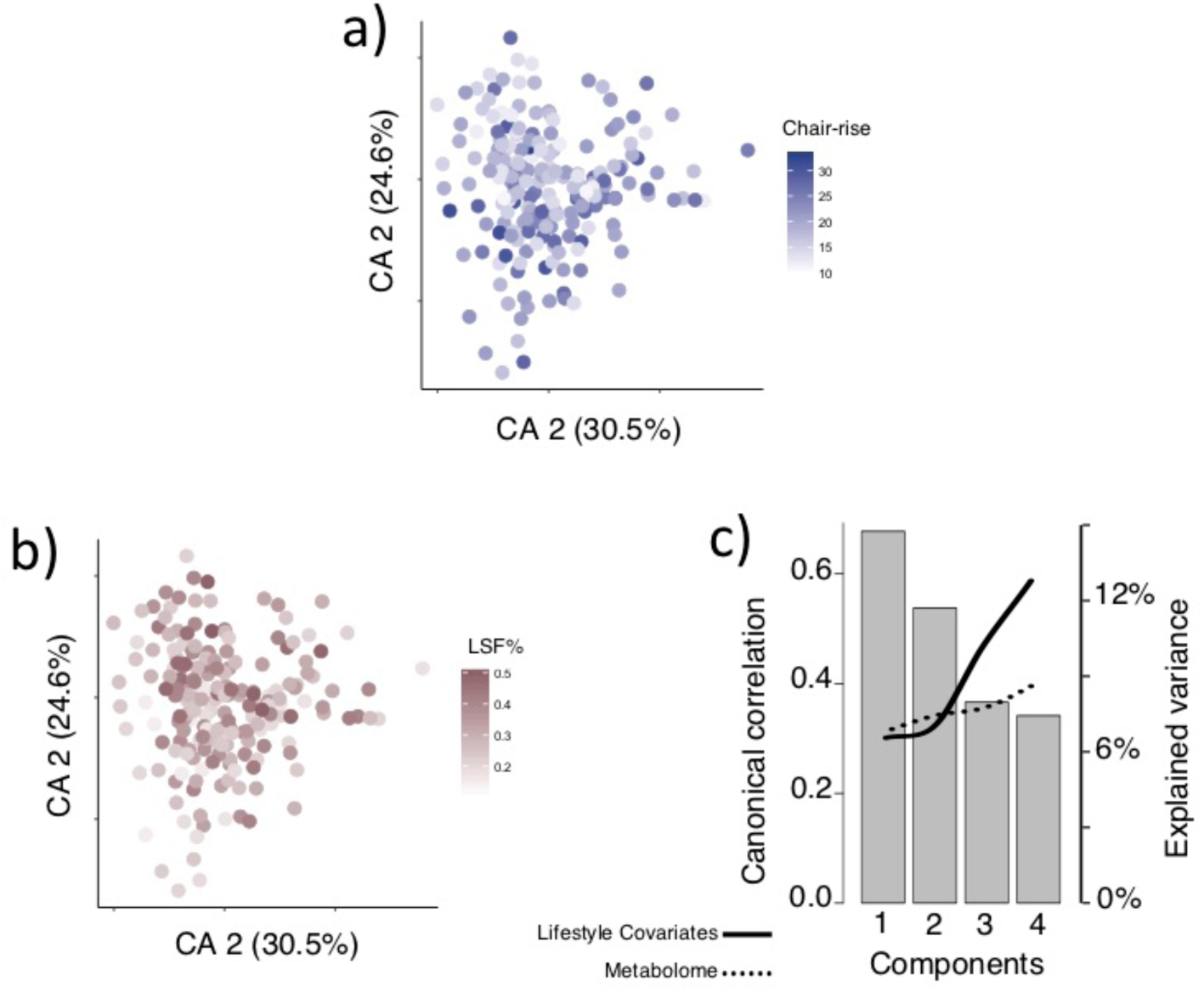
Metabolome correspondence and correlation. (a) Correspondence Analysis of metabolome in relation to chair-stand test and (b) LSF% (c) Canonical correlation within 1^st^ and 4^th^ components as well as explained variance between metabolome profiling and lifestyle covariates.

## Fecal DNA extraction, 16S rRNA-gene amplicon sequencing

Fecal samples were thawed at 4°C, re-suspended in ultrapure water (1:2 feces/water) and homogenized in filter bags for 1 min at high speed (Lab Seward, BA7021). 1.5 ml of the fecal slurry was centrifuged at 13,000×g for 10 min at room temperature and ~200 mg of the fecal pellet was used for DNA extraction using the PowerSoil® DNA Isolation Kit (MOBIO Laboratories, Carlsbad, CA, USA), basically following the instructions of the manufacturer, but with minor modifications to increase lysis of bacterial cells: prior DNA extraction, samples were placed into the PowerBead tubes and heat treated at 65°C for 10 min and then at 95°C for 10 min. Subsequently, solution C1 was added and bead-beating performed in FastPrep (MP Biomedicals, Santa Ana, CA, USA) using 3 cycles of 15 s each, at a speed of 6.5 m s^−1^. The remaining DNA extraction procedure followed the manufacturer’s instructions. Gut prokaryotic composition was determined by NexSeq 500 based 16S rRNA gene-amplicon sequencing of the V3 region amplified using primers designed with adapters for the Nextera Index Kit® (Illumina, CA, USA): NXt_338_F: 5’- TCG TCG GCA GCG TCA GAT GTG TAT AAG AGA CAG ACW CCT ACG GGW GGC AGC AG -3’ and NXt_518_R: 5’- GTC TCG TGG GCT CGG AGA TGT GTA TAA GAG ACA GAT TAC CGC GGC TGC TGG -3’. Amplification profile (1^st^ PCR), barcoding (2^nd^ PCR), amplicon library purification and sequencing were performed as previously described (Pyndt Jørgensen et al. 2014).

## Analysis of high-throughput amplicon sequencing

The raw dataset containing pair-ended reads with corresponding quality scores were merged and trimmed using the following settings, -fastq_minovlen 100, -fastq_maxee 2.0, - fastq_truncal 4, -fastq_minlen 130. Finding unique reads and deconvoluting from chimeric reads and constructing *de-novo* zero-radius Operational Taxonomic Units (zOTU) was conducted using the UNOISE pipeline (Edgar 2018) coupled to the EZtaxon 16S rRNA gene collection as a reference database (Kim et al. 2012). Downstream analyses were based on a contingency table rarefied to 17,000 sequences per sample and then normalized with cumulative sum scaling (CSS (Paulson et al. 2013)).

## Metabolomics

### Untargeted metabolomics of fecal slurries

1 ml fecal homogenate (as described above) was mixed with 1 ml of Sterile PBS (5.7 mM Na2HPO4, 24.3 mM NaH2PO4, 450 mM NaCl, pH 7.4), frozen in liquid nitrogen and freeze-dried overnight. Twenty mg of each freeze-dried sample were re-suspended in 1 ml of 99.98% methanol (containing 10 ppm palmitic-acid methyl ester and 10 ppm sorbitol as an internal standards), vortexed and centrifuged for 30 min at 12,000×g at 4°C. Fifty µl of the supernatant were then dried using a ScanVac (Labogene, Lynge, Denmark) at 1,000 rpm for 3 h at 40°C. Immediately after drying, samples were sealed with air tight magnetic lids into 2.0 ml GC-MS vials and derivatized in two steps using a Dual-Rail MultiPurpose Sampler (MPS) (Gerstel, Mülheim an der Ruhr, Germany), (i) addition of 10 µl of MEOX reagent (20 mg ml^−1^ Methoxiamine hydrochloride in dry pyridine) followed by agitation at 45°C for 90 min by mixing at 750 rpm, (ii) addition of 40 µl of TMS reagent, trimethylsilyl cyanide (TMSCN) (Khakimov et al. 2013) followed by agitation at 45°C for 45 min by mixing at 750 rpm. All steps involving sample derivatization and injection were automated using MPS, which was equipped with a sample agitation unit. Immediately after derivatization, 1 μl of the derivatized sample was injected into a cooled injection system (CIS4) (Gerstel, Mülheim an der Ruhr, Germany) port in splitless mode. The septum purge flow and purge flow to split vent at 2.5 min after injection were set to 25 and 15 ml min^−1^, respectively. Initial temperature of the CIS4 port was 45°C, and heated at 12°C s^−1^ to 320°C (after 30 s of equilibrium time), where it was kept for 10 min. After heating, the CIS4 port was gradually cooled to 250°C at 5°C s^−1^, and this temperature was kept constant during the run. The GC-TOF-MS setup was made combining an Agilent 7890B gas chromatograph (GC) (Agilent Technologies, California, USA) with a time-of-flight mass spectrometer, HT Pegasus TOF-MS, (LECO Corporation, Saint Joseph, USA). GC separation was performed on a Zebron ZB 5% Phenyl 95% Dimethylpolysiloxane column (30 m with I.D. 250 μm and film thickness 0.25 μm) with a 5 m inactive guard column (Phenomenex, Torrance, USA). A hydrogen generator, Precision Hydrogen Trace 500 (Peak Scientific Instruments Ltd, Inchinnan, UK) was used to supply a carrier gas at a constant column flow rate of 1.0 ml min^−1^. The initial temperature of the GC oven was set to 40°C and held for 2 min followed by heating at 10°C min^−1^ to 320°C and kept for an additional 6 min, making the total run time 36 min. Mass spectra was recorded in the range of 45–600 m/z with a scanning frequency of 10 scans sec^−1^, and the MS detector and ion source was switched off during the first 6.3 min of solvent delay time. The transfer line and ion source temperature were set to 280°C and 250°C, respectively. The mass spectrometer was tuned according to manufacturer’s recommendation using perfluorotributylamine (PFTBA). MPS and GC-TOF-MS were controlled using vendor software Maestro (Gerstel, Mülheim an der Ruhr, Germany) and ChromaTOF (LECO Corporation, Saint Joseph, USA), respectively. Samples were randomized prior to derivatization and GC–MS analysis. In order to monitor instrument performance, a blank sample containing only derivatization reagent, a control sample (a pooled sample), and an alkane mixture standard sample (all even C10-C40 alkanes at 50 mg L^−1^ in hexane) were injected after every 10 real samples.

The raw GC-TOF-MS data was processed using Statistical Compare toolbox of the ChromaTOF software (Version 4.50.8.0) with following settings; the raw data was used without smoothing prior to peak deconvolution, baseline offset was set to 0.8, expected averaged peak width was set to 1.5 sec, signal-to-noise was set to ≥10, peak areas were calculate using deconvoluted mass spectra (DT), common *m/z* ions of derivatization products were determined as 73, 75, and 147, deconvoluted mass spectra were also used for peak identification using LECO-Fiehn and NIST11 libraries. The library search was set to return top 10 hits with EI-MS match of >75% using normal-forward search and with a mass threshold of 20. Deconvoluted peaks were aligned across all samples using following settings; retention time shift allowance of <3 sec, EI-MS match of >95%, mass threshold of >25, and present in >90% of all pooled control samples.

### Targeted analysis of SCFA and O/B-CFA in fecal slurries

Analysis of SCFA and O/B-CFA was performed on 0.5 ml of fecal homogenate mixed with 1 ml of 0.3M oxalic acid (containing 2 mM of 2 ethylbutyrate (Sigma-Aldrich) as the internal standard). Samples were vortexed for 1 min, centrifuged at 20°C for 20 min at 12,000×g, followed by filtration using a 0.45 µm centrifugal filter (Millipore UFC30HV00) and the obtained aliquot was used for GC-MS analysis. The GC-MS consisted of an Agilent 7890A GC and an Agilent 5973 series MSD. GC separation was performed on a Phenomenex Zebron ZB-WAXplus column (30 m × 250 μm × 0.25 μm). A sample volume of 1 μl was injected into a split/splitless inlet at 285°C using split mode at 2:1 split ratio. Septum purge flow and split flow were set to 13 ml min^−1^ and 2 ml min^−1^, respectively. Hydrogen was used as carrier gas, at a constant flow rate of 1.0 ml min^−1^. The GC oven program was as follows: initial temperature 100°C, equilibration time 1.0 min, heat up to 120°C at the rate of 10°C min^−1^, hold for 5 min, then heat at the rate of 40°C min^−1^ until 230°C and hold for 2 min. Mass spectra were recorded in Selected Ion Monitoring (SIM) mode and m/z ions were detected at the dwell time of 50 msec: 41, 43, 45, 57, 60, 73, 74, 84. The detector was switched off during the 1 min of solvent delay time. The transfer line, ion source and quadrupole temperatures were set to 230, 230 and 150°C, respectively. The mass spectrometer was tuned according to manufacturer’s recommendation using perfluorotributylamine (PFTBA). Dilution series of SCFA standards of acetic, propionic, butyric, isobutyric, 2-methyl isobutyric, valeric and isovaleric acid (Sigma-Aldrich) were prepared in concentrations of 1.000, 0.500, 0.250, 0.125, 0.060 and 0.030 mM for the construction of standard curves for quantification. Initial inspection of the GC-MS data was performed using MSD ChemStation software (Version E.02.02.1431, Agilent Technologies, Inc., Germany). Mass spectra of SCFA were compared against the NIST11 library (NIST, Maryland, USA). SCFA peak areas were integrated from SIM chromatograms using in-house Matlab (Version. R2015a, The MathWorks, Inc., Massachusetts, USA) scripts. Two SCFA, 2-methyl isobutyric acid and isovaleric acid, co-eluted at the retention time range of 4.22-4.45 min, thus peak areas were calculated by deconvoluting these peaks using *m/z* ions 74 for 2-methyl isobutyric acid and 60 for isovaleric acid.

### Untargeted metabolomics of blood plasma

A mixture of 100 µl of plasma samples (thawed at room temperature) and 300 µl of MeOH:water (8:1, vol:vol and containing 10 ppm of sorbitol as internal standard) were vortexed (highest speed) for 1 min. Thereafter, samples were incubated at 4°C for 15 min and centrifuged at 16,000×g at 4°C for 10 min. Supernatants were passed through a 0.45 µm centrifugal filter (Millipore UFC30HV00) and 80 μl aliquots were dried into 200 μl glass inserts using a ScanVac (Labogene, Lynge, Denmark) at 40°C for 3 h at 1,000 rpm. Immediately after drying samples were sealed with air tight magnetic lids into 2.0 ml GC-MS vials and derivatized in two steps using MPS, (i) addition of 10 µl of MEOX reagent (20 mg ml-1 Methoxiamine hydrochloride in dry pyridine) followed by agitation at 65°C for 60 min by mixing at 750 rpm, (ii) addition of 30 µl of TMS reagent (TMSCN) followed by agitation at 65°C for 2 h by mixing at 750 rpm. Immediately after derivatization, 1 μl of the derivatized sample was injected into the GC-TOF-MS as described for the fecal metabolomics. Sample injection, oven and mass spectrometer parameters were similar to those for the fecal metabolomics with few modifications. The initial temperature of the GC oven was set to 40°C and held for 2 min followed by heating at 12 °C min^−1^ to 260°C, and with a rate of 30°C min^−1^ to 320°C and kept for an additional 5 min, making the total run time 27.33 min. Mass spectra was recorded in the range of 45–500 m/z with a scanning frequency of 8 scans sec^−1^, and the MS detector and ion source was switched off during the first 8.3 min of solvent delay time. The transfer line and ion source temperature were set to 290°C and 250°C, respectively. In order to monitor instrument performance, a blank sample containing only derivatization reagent, a control sample (a pooled sample), and an alkane mixture standard sample (all even C10-C40 alkanes at 50 mg L^−1^ in hexane) were injected after every 10 real samples. The raw GC-TOF-MS data was processed as described above for untargeted fecal metabolomics.

